# Diversity, function and evolution of marine microbe genomes

**DOI:** 10.1101/2021.10.26.465843

**Authors:** Jianwei Chen, Yang Guo, Yangyang Jia, Guilin Liu, Denghui Li, Dayou Xu, Bing Wang, Li Zhou, Ling Peng, Fang Zhao, Yuanfang Zhu, Jiahui Sun, Chen Ye, Jun Wang, He Zhang, Shanshan Liu, Inge Seim, Xin Liu, Xun Xu, Huanming Yang, GOMP Consortium, Karsten Kristiansen, Guangyi Fan

## Abstract

Trillions of marine bacterial, archaeal and viral species contribute to the majority diversity of life on Earth. In the current study, we have done a comprehensive review of all the published studies of marine microbiome by re-analyzing most of the available high throughput sequencing data. We collected 17.59 Tb sequencing data from 8,165 metagenomic and prokaryotic samples, and systematically evaluated the genome characters, including genome size, GC content, phylogeny, and the functional and ecological roles of several typical phyla. A genome catalogue of 9,070 high quality genomes and a gene catalogue including 156,209,709 genes were constructed, representing the most integrate marine prokaryotic datasets till now. The genome size of Alphaproteobacteria and Actinobacteria was significant correlated to their GC content. A total of 44,322 biosynthetic gene clusters distributed in 53 types were detected from the reconstructed marine prokaryotic genome catalogue. Phylogenetic annotation of the 8,380 bacterial and 690 archaeal species revealed that most of the known bacterial phyla (99/111), including 62 classes and 181 orders, and four extra unclassified genomes from two candidate novel phyla were detected. In addition, taxonomically unclassified species represented a substantial fraction of 64.56% and 80.29% of the phylogenetic diversity of Bacteria and Archaea respectively. The genomic and ecological features of three groups of Cyanobacteria, luminous bacteria and methane-metabolizing archaea, including inhabitant preference, geolocation distribution and others were through discussed. Our database provides a comprehensive resource for marine microbiome, which would be a valuable reference for studies of marine life origination and evolution, ecology monitor and protection, bioactive compound development.

## Introduction

Marine microbes, which includes viroids, viruses, bacteria, archaea, fungi and protists varies from non-cellular viruses, single-cell organisms to multicellular microorganisms, encompassing all three domains of life. About 10^30^ prokaryotes cells and ten-times more femtoplankton (viruses) are estimated in the oceans, comprising the majority of global microbial biomass and 90% of ocean biomass[1–3]. After 3.5 billion years of evolution, microbes account for the major fraction of the marine biodiversity, abundance, and metabolism, and play fundamental roles in sustaining the development and maintenance of all other marine lives and their activities [1, 4]. The enormous and highly diverse marine microbes are responsible for up to 98% of primary marine productivity in global cycling of nutrients, matter, and energy in the oceans through biogeochemical processes (carbon, nitrogen, sulfur cycling, etc.) [3, 5, 6]. Furthermore, marine microbes produce a plethora of natural biologically active products with such as cytotoxic, antifoulants, anti-inflammatory, anti-viral, antifungal, antibacterial and anti-tumor activities [7, 8], which represent important and promising sources for new drug discovery and drug development [8–10].

While in terrestrial ecosystem, higher plants work as the main group of primary producers, it is prokaryotes and other microbes who play that role in marine ecosystem[3]. What is more, marine prokaryotes have been demonstrated to regulate the biogeochemical cycles and the climates on a large scale, such as the global carbon cycle [11–13], nitrogen cycle [14, 15], and green-house effect [16, 17]. For example, in the case of carbon cycle, both phototrophic and chemoautotrophic marine prokaryotes as well as other organisms such as algae and protists using light or chemical energy to fix carbon into cellular material [18], among which, Cyanobacteria are recognized as main contributors [19–21]. Occupying a broad range of habitats across all latitudes, and even the most extreme niches [22], Cyanobacteria absorb more than 2/3 of the total carbon sequestration in the ocean each year [23]. However, on the other hand, the dense cyanobacteria blooms which sometimes are toxic could threaten ecosystem and human health [24].

In addition to carbon sequestration, marine sediment methane and hydrates, accounting for the vast majority of methane pool on the earth, represent another major form of carbon in the ocean. However, only quite a small fraction of the seabed methane could be released to the atmosphere[25, 26]. In marine sediments, the biogenic methane is exclusively produced by methanogenic archaea in strictly anaerobic environments[27], meanwhile, Ca. 80∼90% of the global methane gross production from marine sediments is oxidized by methanotrophic microbial communities [26]. Thus, microbes of both methanogens and methanotrophs exert a major control on global climate and carbon (C) cycles, since methane could cause 25 times of green-house effect compared to CO_2_ [28].

Marine prokaryotes are also closely related to human beings. Such as some *Vibrio* bioluminescence are useful as a biomarker during scientific experiments, and provides abundant bioactive substances including medicines and cosmetics[29–31]. Except for economical and medical product derived from them, many marine prokaryotes are potential pathogens to human, which is one of the major threats of heath especially for people working in shipping and fishery industry [32, 33]. For example, many *Vibrio* species are pathogenic[34], and marine *Vibrio* species can infect human with interaction on coastal biomes[35].

Despite the global importance of marine prokaryotes, most of them remain untouched and thus still are “dark matter” till now, either due to being unculturable or their extreme diversity or rarity for some taxa[36, 37]. High-throughput sequencing techniques now allow us to quickly obtain genome sequences of theoretically all the species in certain environments without culturing. The metagenomics sequencing has become an important tool for studying the composition of microorganisms in various marine ecosystems, such as free-living bacterioplankton[38, 39], the sediment-dwelling microbes[40, 41] and animal-associated symbionts[42, 43]. The Global Ocean Sampling Expedition (GOS) and Tara Ocean Expedition increased our understanding of marine microbial diversity and genetic characteristics vastly [44, 45]. However, the genome sequencing and data mining of marine microbiome are still challenging, as revealed by the slowly increased genome sequence of marine prokaryotes in public database [46]. There are more than 280,000 prokaryotic genomes in public databases, but only 8,615 marine prokaryotic genomes were found. Although many research efforts have been devoted to the marine microbial study and great amount of sequencing data have been generated till now, there is not a comprehensive summary of the previous work, neither a good representative database that could be use as marine prokaryotes genome reference catalogue. And it investigated that metagenomics and bioinformatics are the powerful tools for massive expansion knowledge of microbial genomics research [47, 48].

Thus, in this study, we comprehensively collected and analyzed all the publicly available marine metagenomic high-throughput sequencing data from NCBI and EBI. After re-analyze all those data, we generated a marine prokaryotic genome catalogue included more than 20,000 genomes belonging to 113 phyla, and describe the massive diversity and globally distribution of marine prokaryotes. The discovery of a large number of novel species has expanded the understanding of marine microbial diversity. In addition to that, we also illustrated the main functions of marine prokaryotes in various ocean ecosystems. Our resource will provide new foundation for studies about how the marine microbes adapt to varying environmental conditions and how the marine microbes affect the function and health of marine ecosystems. Furthermore, the attractive genome-based mining of biosynthetic gene clusters (BGCs) provides new insights for the screening of marine bioactive substances and the synthesis of novel active compounds.

## Results

### Benchmark of data set

Sequencing data or assembled genomes where available, of a total of 8,165 prokaryote genomic or metagenomic samples from the marine ecosystem, including seawater, algae and marine animal symbiotic microbiome, mangrove and marine sediment were downloaded from public databases. This dataset covered a broad range of the entire ocean across the earth, with 3,089 samples isolated from Pacific Ocean, 1,396 from Atlantic Ocean, 599 from Indian Ocean, 128 from Arctic Ocean, and 123 from Southern Ocean (**Fig. 1a**). And then all data was used to generate the marine prokaryotic genome and protein sequence catalogs (**Fig. S1**). This is the most comprehensive survey and summary of the microbiome and their genome function and diversity in global marine ecosystems to date. Firstly, the genomes of 10,598 isolate prokaryotic strains or metagenomics assembled genomes (MAGs) were downloaded. Among the 10,598 genomes, 8,300 of them were moderate genomes (completeness >50%, contamination <10%), of which 6,213 were substantial genomes (completeness >70%, contamination <10%), and of the substantial genomes, 4,629 were near complete genomes (completeness >90%, contamination <5%). In the current study, only the 6,213 substantial genomes were selected and retained for downstream analysis (**Fig. 1b**). Meanwhile, more than 17.59 Tb sequencing data of 2,695 samples were used for assembly and binning analysis respectively. A total of 20,671 moderate prokaryotic MAGs including 14,969 substantial MAGs were reconstructed, and only the 14,969 substantial genomes including 5,938 near complete genomes were remained for downstream analysis as well (**Fig. 1b**). Besides, in the unique gene catalogue we constructed, a set of 156,209,709 genes were included, which was near four times larger than the Tara Ocean gene set [44].

**Fig.1.**
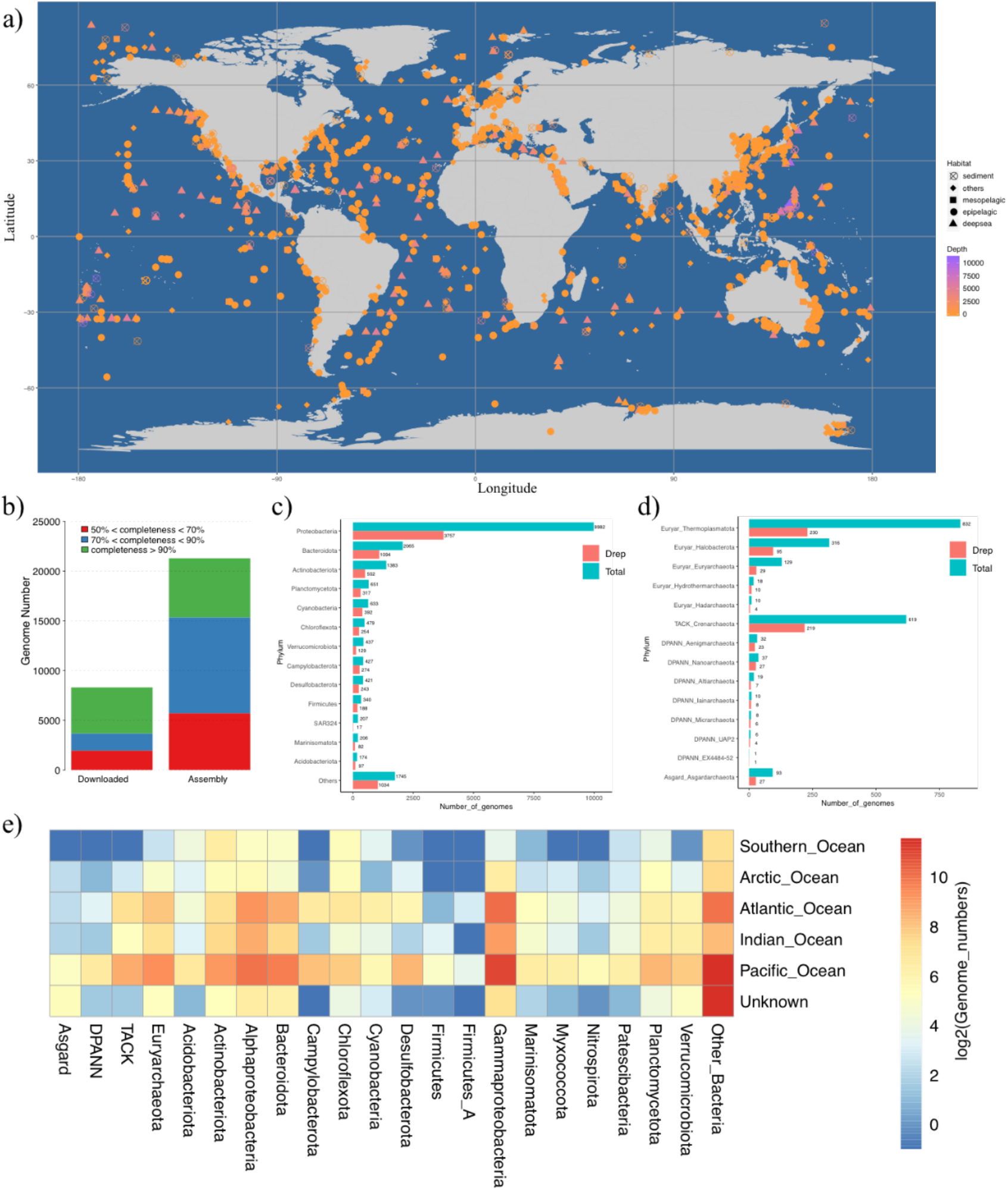
Benchmark of the data set. a) Distribution of the samples collected in the current study. Summary of the quality of the genomes with contamination <10% b), and taxonomic annotation of the assembled genomes at phylum level for bacteria c) and archaea d). e) The genome distribution among the different Ocean regions.

After taxonomic classification for all downloaded genomes and assembled MAGs, 21,182 high quality prokaryotic genomes including 19,064 bacterial genomes and 2,118 archaeal genomes were obtained (**Fig. 1b**). And we generated a unique species-specific genome catalogue of the marine microbiome basis 95% nucleotide identity threshold, including 8,380 unique bacterial genomes and 690 unique archaeal genomes while only 3,753 genomes were from public database. The genome catalogue generated in our study greatly exceed the previous results, such as 2,631 moderate genomes including 420 near complete genomes generated from 243 Tara Ocean microbial metagenomic samples[44, 49], and 4,741 and 8,578 moderate genomes generated by GORG-Tropics Database [50] and Earth’s Microbiomes Project [51], respectively. We detected 97 bacteria phyla, with Gammaproteobacteria, Alphaproteobacteria, Bacteroidota, Actinobacteriota, Planctomycetota and Cyanobacteriia being the most common phyla, containing 5,875, 4,201, 2,114, 1,408, 665 and 643 assembled genomes respectively, all of which are the most common bacterial populations (**Fig. 1c**). In addition to the previous defined 97 bacterial phyla, two novel bacterial phyla were detected and annotated by GTDB-tk, and here we name them as candidate phylum MSD20-3 and candidate phylum MSD20-1. We also obtained 14 archaea phyla are detected with Euryarchaeota and TACK being the mainly assembled archaeal genomes (**Fig. 1d**).

We further studied the global distribution of the marine microbes, and found that the prokaryotic species distribution is quite different in different marine ecological systems. For example, the bacterial species in different marine habitats, including coastal surface waters, open seas, and sediments are very different from each other. There are about 57.90% samples distributed in Pacific Ocean, and we found that more than 61.74% archaeal genomes and 56.56% bacterial genomes were detected in this ocean (**Fig. 1e**). Archaeal species rarely detected in the Southern Ocean, with only six Euryarchaeota genomes detected in Antarctic Ocean. The Actinobacteriota, Chloroflexota and Gammaproteobacteria are the common species in polar regions, while Gammaproteobacteria, Alphaproteobacteria and Bacteroidota are the top three abundant species distributed in Atlantic Ocean, Indian Ocean and Pacific Ocean.

### Phylogenetic evolution of marine bacteria and marine archaea

The phylogenetic distribution of the 8,380 bacterial (**Fig. 2a**) and 690 archaeal (**Fig. 2b**) species revealed that taxonomically unclassified species represented 64.56% and 80.29% of the phylogenetic diversity of Bacteria and Archaea respectively. However, previously only 13 bacterial phyla with 22.55% unclassified species genomes and two archaeal phyla (Euryarchaeota and Thaumarchaeota) with 18.25% unclassified species genomes were found in Tara Ocean MAGs[49]. The large fraction of the unclassified genomes indicates that there are still many prokaryotes that have not been studied in the marine ecosystem. Most bacterial phyla (99/111) were detected and two newly phyla included four genomes, 62 classes unclassified genomes and 181 orders unclassified genomes were reported (**Fig. 2a &2c**). The first new phylum candidate phylum MSD20-3 was phylogenetically close to phylum Elusimicrobiota, and three draft genomes retrieved from SRR11637895 (bin.20), SRR9661844 (bin.98) and SAMN10404973 (bin.31) were included (**Fig. 2a**), while the second new phylum candidate phylum MSD20-1, including one draft genomes retrieved from SAMN1451138 (bin.12), was phylogenetically close to phylum Hydrogenedentota (**Fig. 2a**). The average nucleotide identity (ANI) between the new phyla and their respective most phylogenetically close relatives are both ∼60%, indicating large divergence distance between the genome of new phyla and their close relatives [52].

**Fig. 2.**
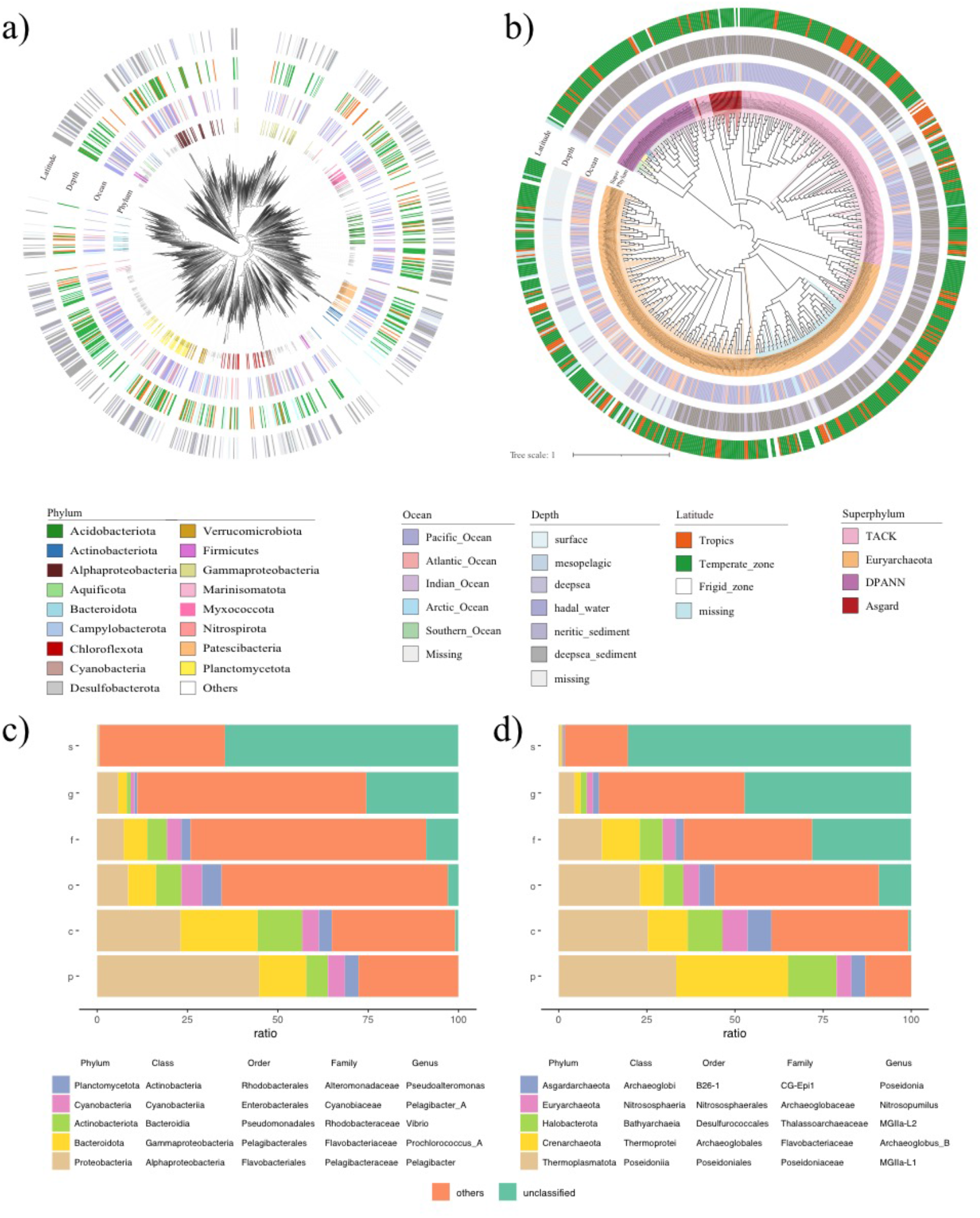
Phylogenetic tree and the proportion of different level of marine bacteria and archaeal genome. The phylogenetic tree and sample metadata of 8,380 marine bacteria genomes a) and 690 marine archaea genomes b). The top five abundant species and unclassified genomes in different taxonomic levels of bacteria c) and archaea d).

Compared with bacteria, our knowledge of archaea is still very limited. In previous studies, microbiologists explore archaea mainly by means of pure culture or single-gene diversity survey. However, only 22% known archaea phyla have isolated and cultured representative species [53]. Here, we constructed 690 archaeal genomes distributing in 14 archaeal phyla (total 18 phyla), and five class unclassified genomes, 58 order unclassified genomes were firstly found with a high unclassified species proportion (**Fig. 2b&2d**). Among the 690 unique archaea genomes, Euryarchaeota takes up the highest proportion (56.7%), followed by TACK (31.7%) and DPANN (7.7%), Asgard (3.9%) has the least proportion (**Fig. 2b**). Especially we constructed 93 high quality Asgard archaea genomes and obtained 27 de-redundant genomes included one unclassified class. It was helpful for refining the phylogenetic relationships of Asgard and adding new evidence of the earliest evolutionary history of life [54].

### Genome features of marine prokaryotes

The genome size and GC content vary greatly in different marine bacteria. The genome size of most marine bacteria ranges from 2Mb to 5Mb, harboring mostly 3000-5000 genes, with GC content ranging from 30% to 60% (**Fig. 3a**). However, for bacteria in certain phylum, they have extraordinary genome features. For example, Patescibacteria has the smallest genomes with an average of only 0.80 Mb, followed by Aquificota with averaged genome size of 1.37 Mb (**Fig. 3a**), while the largest genomes belong to Myxococcota phylum, with an averaged genome size of 5.84 Mb. Likewise, for the GC content of marine bacteria, Firmicutes_A has the lowest GC content of 33.34%, while Myxococcota has the highest GC content of 63.47% in average (**Fig. 3a**).

**Fig. 3.**
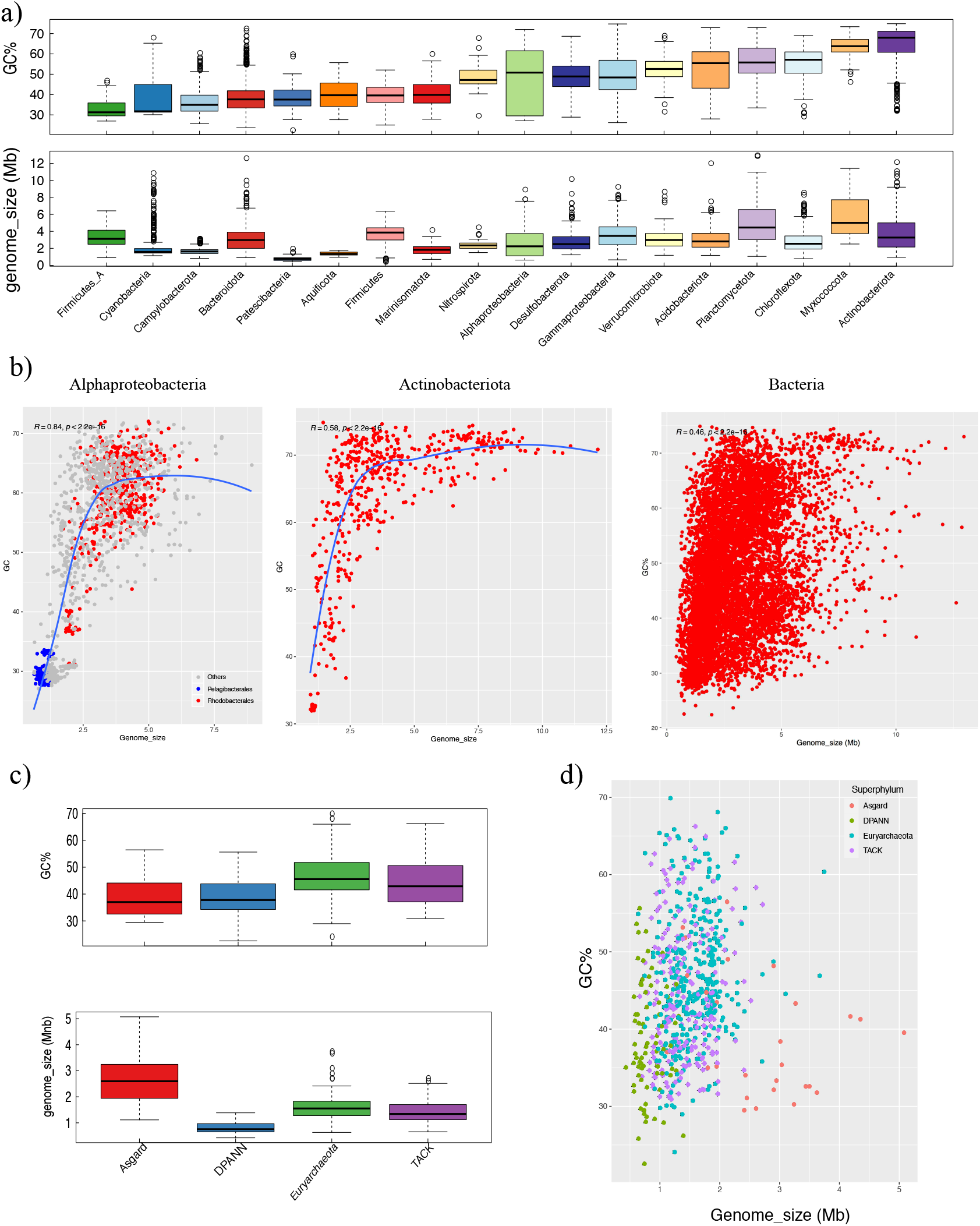
Summary and comparison of the genome size, GC content of marine bacteria and archaea genome. The genome size and GC content statistics of major bacteria group a) and archaea superphylum c). And the genome size and GC content correlation of Alphaproteobacteria, Actinobacteria and all marine bacteria b) and all marine archaea d).

Spearman correlation analysis indicates that the genome size and GC content of marine bacteria has an overall significant positive correlation (R=0.46, P<2.2e-16). The correlation coefficient between genome size and GC content of Alphaproteobacteria (R=0.84, P<2.2e-16) and Actinobacteria (R=0.58, P<2.2e-16) are even higher than the overall correlation coefficient (**Fig. 3b**). However, despite the significant positive correlation, we found that as the genome size increased, the GC content of these two species increased at first and finally reached the upper limit of 75%, which is in accordance with the GC compositional range of prokaryotes between approximately 25% and 75% [55]. Furthermore, the GC content as a function of genome size distribution is not linear but triangular, to which similar distribution pattern was also observed in previous studies of bacteria[56], vertebrates[57] and plants[58].

In the phylum of Alphaproteobacteria, species with small size and low GC content (GC<35% and genome size<2.5M) were Pelagibacterales (1,017 of 1,274 genomes, blue), HIMB59 named Pelagibacteraceae (159 genomes) and Puniceispirillales (26 genomes) as colored in blue at the left bottom of **Fig. 3b** (**Fig. 3b**). Pelagibacterales (SAR11) are one of the smallest free cell living organisms (<0.7 um) composed of free-living planktonic oligotrophic facultative photochemotroph bacteria[59]. Their high surface to volume ratio guarantees them better capability to absorb nutrients from its oligotrophic environment, and oxidize organic compounds from primary production into CO_2_ [60]. The species dominant in the top right of Alphaproteobacteria (GC>35% or genome size>2.5M) were Rhodobacterales (1042 of 2872 genomes, red), Sphingomonadales (410 genomes), and Caulobacterales (357 genomes) (**Fig. 3b)**. Rhodobacterales are widespread in the marine ecosystem and show a nearly universal conservation of the genes for production of gene transfer agents (GTAs) which are virus-like particles[61]. Thus, our result indicated that transfer DNA might mediate genetic exchange between cells and be an important factor in their evolution.

We found the GC frequency of the third base of the codon is very low (only 18.31%) in the Pelagibacterales genomes with low GC content and small genome size (**Table 1**). For the Rhodobacterales with a wide GC distribution and genome size, the species with larger genome size (>2.5 Mb) have higher GC content than the species with smaller genome size (<2.5 Mb), and the third-base GC frequency of the codon is significantly higher (**Fig. 3b, Table 1**). And we have also observed the consistent patterns in Actinobacteria (**Table 1**). It indicated that in high GC species, the third base of the gene codons with higher variability is more inclined to use the G+C base instead of A+T base.

**Table 1.**
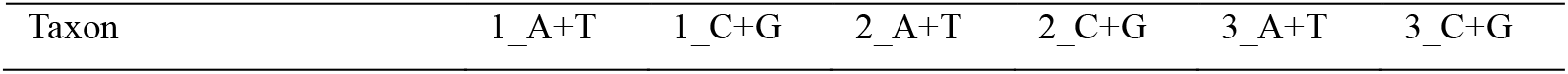
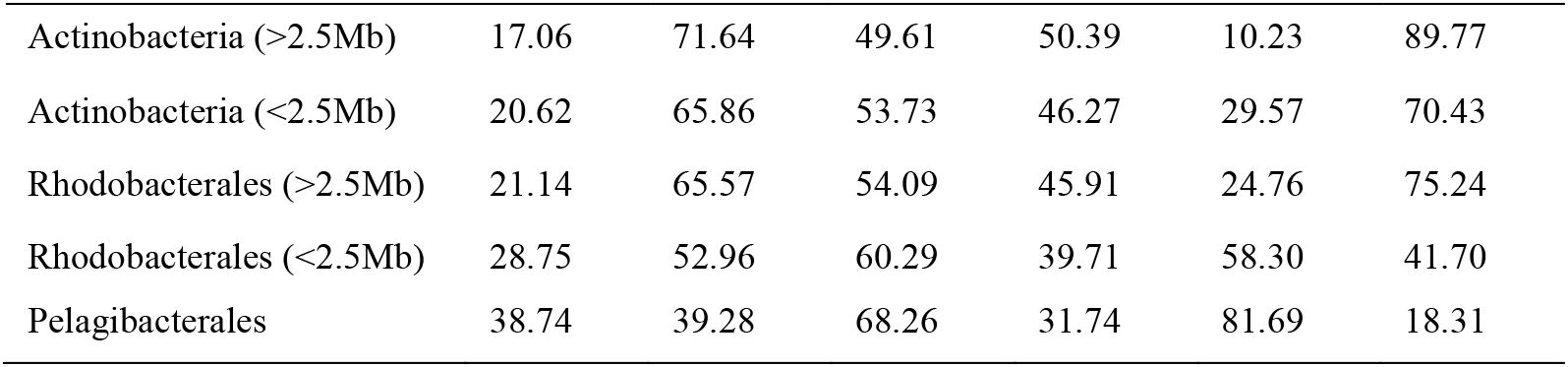
Codon base frequency statistics.

For the marine archaeal genomes, the GC ratio is ranging from 30% to 55% with genome sizes of ∼1-3Mb (**Fig. 3c & 3d**). No significant correlation between genome size and GC content in the marine archaea genomes was found (**Fig. 3d**). Most genomes belonging to DPANN superphylum have extremely small cell and genome sizes (∼0.5 to 1.5 Mb, averaged 0.82M) with limited metabolic capabilities [62]. The DPANN genomes also have lowest GC content (averaged 38.60%) in archaea genomes. For example, MAG SRR5506558.1_bin.59 (completeness 72.9%) has the smallest genome size of 0.42M with 35.04% GC content, which is smaller than the previously reported *N*. *equitans* (GCA_000008085.1) with genome size of 0.49 Mb and completeness 73.13%[63]. SRR5214304_bin.64 (completeness 89.72%) has the largest genome size of 1.75M in DPANN superphylum with GC content of 38.77%. In archaea kingdom, genomes in the phylum of Asgard have the largest genome size (average 2.68M), which indicates more complex genome structure and content than other marine archaea.

As in other environments, the genome size, GC content and distribution of microbes are related to and restricted by physiochemical and nutritional conditions in marine environments. Consistent with previous reports, most bacterioplankton and pelagic dwelling bacteria, including Pelagibacterales (SAR11), Synechococcus, Prochlorococcus and Thioglobaceae (SUP05) usually have low GC content (∼28-40%) and small genome size (∼0.8-3Mb) (**Fig. 4a**) [64]. In contrast, both the GC content (33-73%) and genome size (1-13Mb) of Myxococcota, Planctomycetota and archaea Euryarchaeota ranged widely and distributed from the surface ocean to deep-sea. The Puniceispirillaceae (SAR116), Patescibacteria and archaea TACK superphylum have small genome size but widely ranging GC content, of which while the Puniceispirillaceae is surface dwelling and the other two are living in various depth ocean layers (**Fig. 4a**). The Archaea clades Euryarchaeota, TACK superphylum and Bacteria clades Planctomycetota, Patescibacteria, Myxococcota occupy extreme low temperature environments distributing from the Antarctic to the Arctic, while most species of Puniceispirillaceae, Pelagibacterales, Synechococcus, Prochlorococcus and Thioglobaceae thrive in the temperate zone with small genome size (**Fig. 4b**).

**Fig. 4.**
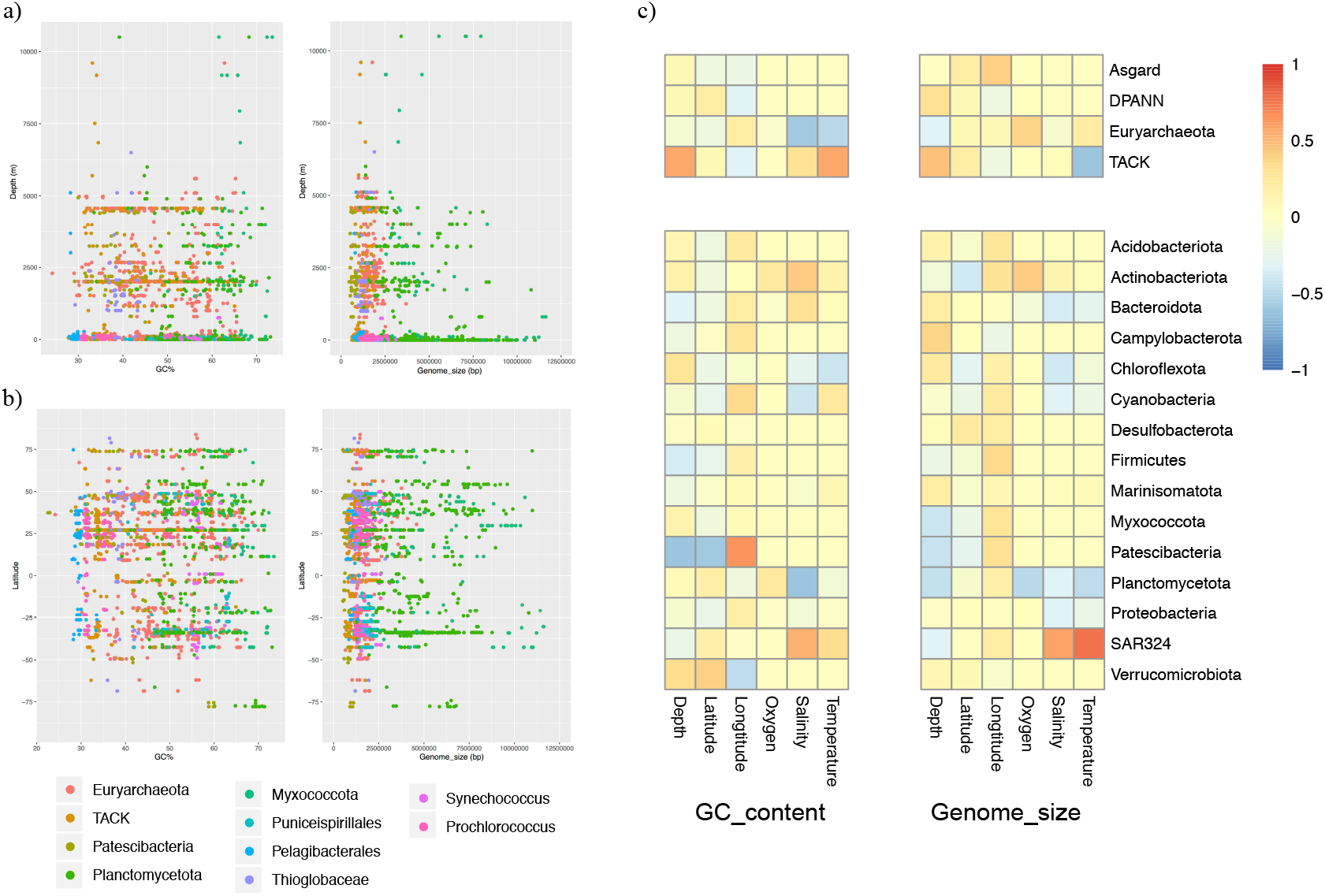
Correlation analysis between the environmental factors and GC content and genome size. The distribution of 10 major species at different depth a) and latitude b). c) The correlation analysis heatmap of environmental factors and GC content and genome size.

Correlation test between marine microbial genome size and GC content with various environmental factors were conducted using spearman correlation analysis. The GC content of Euryarchaeota and Planctomycetota decreased significantly with salinity, while the GC content of Patescibacteria decreased significantly with depth and latitude (**Fig. 4c**). The GC content and genome size of SAR324 clade increased significantly with salinity and temperature, and GC content of TACK superphylum increased significantly with depth and temperature while the genome size decreased with temperature (**Fig. 4c**).

### Gene function analysis and BGCs detection

Functional genes predicted from the reconstructed genomes were annotated against the KEGG database. The annotated proportion of functional genes in different marine prokaryotes are quite different. Totally, 240 KEGG pathways were detected in Actinobacteriota genomes, followed by 237 and 218 pathways detected in Gammaproteobacteria and Firmicutes respectively (**Fig. 5a**). Not surprisingly, species with the smaller genome size seem to be annotated with fewer pathways, for example, 131 pathways were annotated in DPANN superphylum and 99 pathways in Patescibacteria, indicating that genome-reduction are accompanied with loss of metabolic functions [62, 65]. Biosynthesis of secondary metabolites (ko01110), Biosynthesis of antibiotics (ko01130) and Biosynthesis of amino acids (ko01230) are the most common pathway and the largest proportion genes in most marine prokaryotes except for the DPANN superphylum and Patescibacteria. In particular, Actinobacteriota, Firmicutes and Cyanobacteria contain an average of more than 270 genes per genome annotated to the pathway of Biosynthesis of secondary metabolites, which indicates that a huge number of potential marine bioactive substances.

**Fig. 5.**
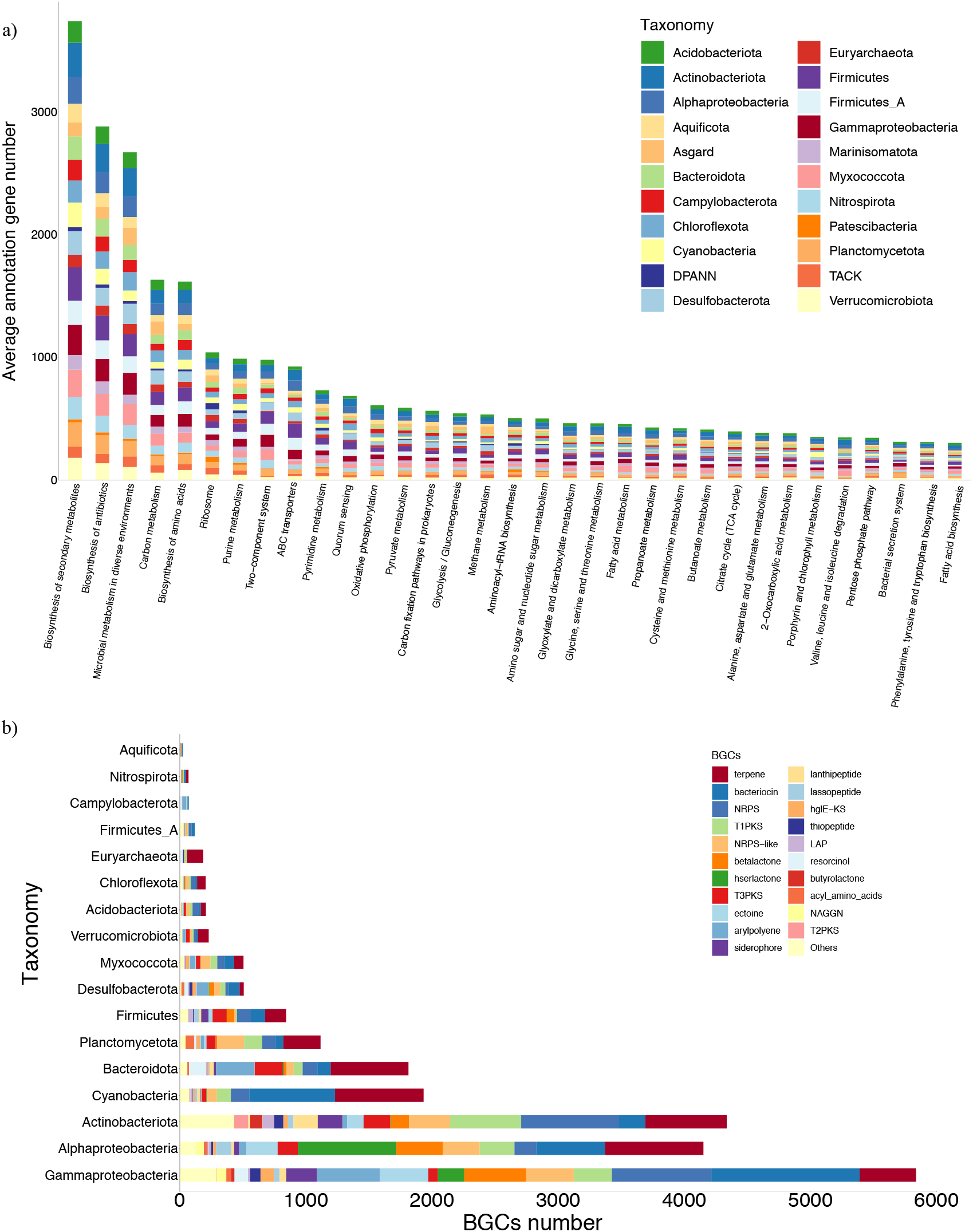
Summary of the KEGG functional annotation and secondary metabolite BGCs.

Meanwhile, we detected more than 53 types of biosynthetic gene clusters (BGCs) in marine bacterial genomes, and predicted 193 BGCs belong to 16 types in marine archaeal genomes (**Fig. 5b**). In archaea, main types of terpene, T1PKS, resorcinol, thiopeptide, TfuA-related, betalactone, bacteriocin and ectoine were found in Euryarchaeota [66], while fewer types of phosphonate, NRPS and T3PKS were found in TACK and Asgard genomes [67].On the other hand, terpene, bacteriocin, NRPS and NRPS-like, T1PKS and T3PKS, arylpolyene and hserlactone are the most common BGCs occur in marine bacteria (**Fig. 5b**). For example, marine Cyanobacteria and Actinobacteriota can produce a wide variety of bioactive substances with various potential functions, such as antibacterial, anti-tumor, anti-virus, cytotoxicity, anti-coagulation and blood pressure reduction. At present, more than 50% of newly discovered marine microbial bioactive metabolites are produced by Actinobacteriota [68]. In the current study, we found 1,101 NRPS and NRPS-like, 646 terpene, 564 T1PKS and 208 bacteriocin BGCs in 502 Actinobacteriota genomes, and 702 terpene, 680 bacteriocin, 224 NRPS and NRPS-like and 111 T1PKS were found in 392 Cyanobacteria genomes.

### Cyanobacteria diversity in marine ecosystem

Due to their extraordinary ability to fix nitrogen and carbon, Cyanobacteria are arguably the most successful group of microorganisms on Earth, playing important roles in the global ecology[69, 70]. They can produce oxygen through photosynthesis system PSI and PSII [71], and fix CO_2_ into organic carbon via ### system [72]. *Prochlorococcus* and *Synechococcus* are the most abundant photosynthetic organism on Earth, especially *Prochlorococcus*, which is responsible for a large fraction of marine photosynthesis.

A totall of 632 Cyanobacteria genomes (388 *Prochlorococcus* and 128 *Synechococcus*), of which 461 were downloaded from NCBI and 171 were newly generated MAGs in the current study. For geographical distribution, 255 Cyanobacteria were distributed in Atlantic Ocean, 224 in Pacific Ocean, 18 in Indian Ocean, 2 in Arctic Ocean and 13 in Southern Ocean. The species *Phormidium* and *Leptolyngbya* are taxonomically unique genotypes and endemic or restricted to polar habitats [73]. And we also found another three species, including Elainellales, Neosynechococcales and Obscuribacterales, specifically distributed in Southern Ocean. Phylogenetic analysis of evolution and geographical distribution indicated that *Prochlorococcus* and *Synechococcus* were clearly separated clades and had no obvious association with the ocean areas, mostly distributed in the ocean area between 40° south latitude and 45° north latitude (**Fig. 6**) [74].

**Fig. 6.**
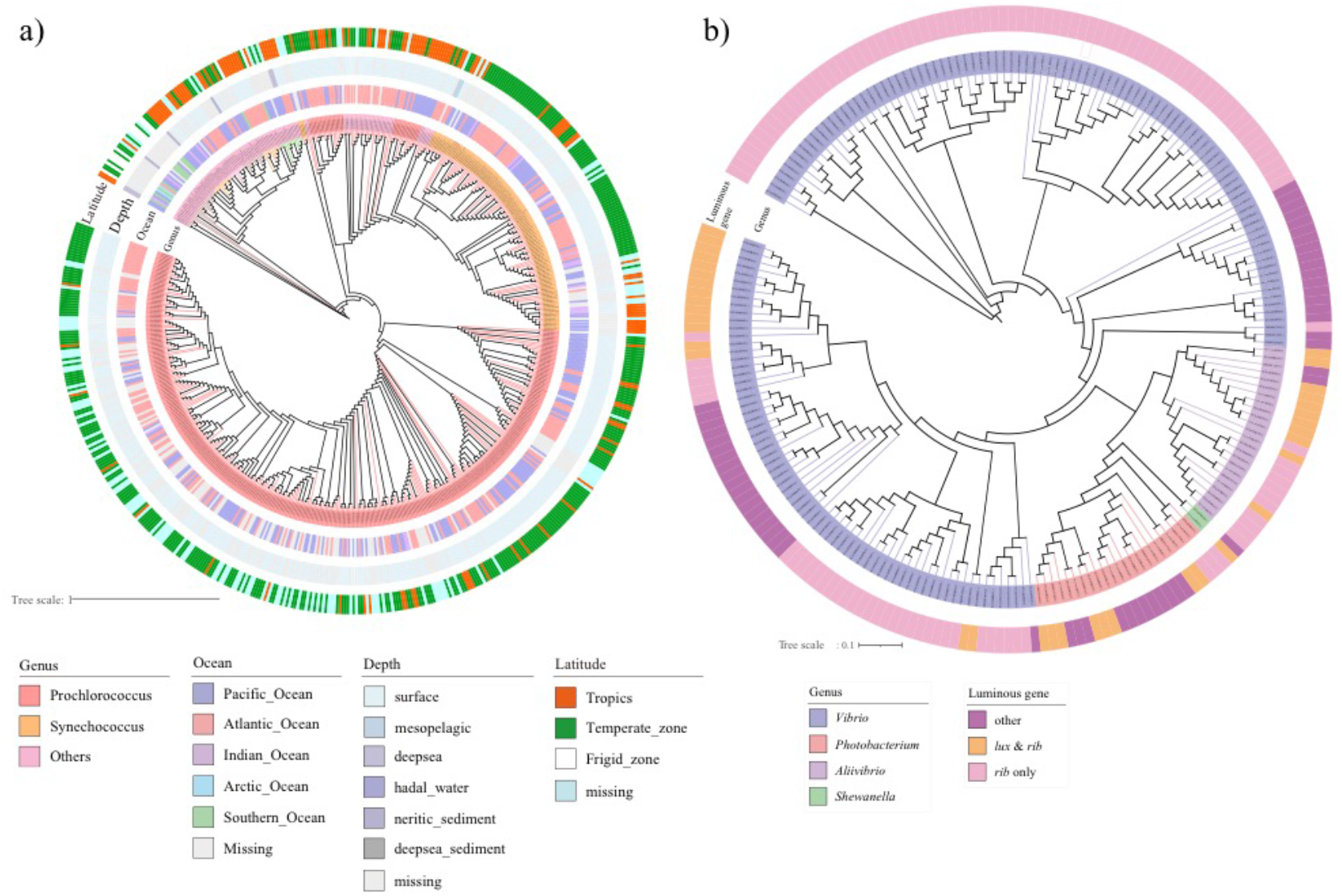
The phylogenetic tree of Cyanobacteria and marine bioluminescent bacteria.

While Cyanobacteria are usually distributed in surface oceans, we reconstructed four high quality Cyanobacteria MAGs (completeness > 80%, contamination < 5%) in the 4000 meters deep-sea of Pacific Ocean, two whichwere classified as *Richelia intracellularis*_B. Meanwhile, four *Richelia intracellularis*_A MAGs were reconstructed in shallow water of 2 to 4 meters of Atlantic Ocean. Thus, we intended to find the difference between deep-sea and shallow water *R*. *intracellularis* genomes. GC content of *R*. *intracellularis*_B MAGs is higher than *R*. *intracellularis*_A, suggesting the huge pressure of the deep ocean may require higher GC content to maintain the stability of the genome [75]. Furthermore, proteins involved in the photosynthesis pathways, such as, the photosystem proteins K02722, K02718, K02712, K02706, K02692 and K02689 were detected in *R*. *intracellularis*_A, while missing in deep-sea *R*. *intracellularis*_B. On the other hand, *R*. *intracellularis*_B contained several unique gene functions related to photosystem II oxygen-evolving enhancer protein and cytochrome including K08904, K02717, K02643 and K08906 which might relate to temperature adaptation[76], all of which were missing in shallow-water *R*. *intracellularis*_A genomes.

### Marine luminous bacteria genome detection

Bioluminescence is a widespread natural phenomenon involving visible light emission, which is advantageous for luminescent organisms through prey luring, courtship display, escaping from predators by dazzling and camouflage via counter illumination [77, 78]. There discovered nearly 800 genera containing thousands of luminescent species, and the vast majority of which reside in the ocean [79, 80]. Although fish and crustaceans are the largest bioluminescent groups by biomass, bacteria dominated in terms of abundance. By far, luminous bacteria have been found among in three families of Vibrionaceae (*Vibrio*, *Photobacterium*, *Aliivibrio* and *Photorhabdus*), Shewanellaceae (*Shewanella*) and Enterobacteriaceae. Except for *Photorhabdus* in the five classified luminous genera, all the ther other four genus, including *Vibrio*, *Photobacterium*, *Aliivibrio* and *Shewanella*, could reside in the sea[81]. Here in the current study, we classified 213 luminous genomes assigned into 164 *Vibrio* (550 *Vibrio* genomes in total), 23 *Photobacterium* (49 genomes in total), 24 *Aliivibrio* (37 genomes in total) and 2 *Shewanella* (41 genomes in total) (**Fig. 5b**). Among of them, one *Alliibrio fischeri* genome could live symbolic or free-living style through the aquatic environments and when could make the animal organs glowing (**Fig. 5b**). In addition, no genome data of luminous Enterobacteriaceae was detected in our genome catalogue.

All luminous bacteria are thought to share the same unique luminescent mechanism. In bacterial luminescent reaction, enzymes encoded by the *lux* operon mediate the oxidation of reduced flavin mononucleotide (FMNH2) produced by *rib* operon and long-chain fatty aldehyde (RCHO) to emit blue-green light[82]. The genetic *lux* operon responsible for luminescence has been well understood. We screened the species and strains has *lux* and *rib* operon (contain genes involved in the synthesis of riboflavin) and found that many luminous Vibrionaceae species or strains apparently lack lux operon, while *lux* operons were detected in some nonluminous species (**Fig. 5b**). It is not clear about the mechanism and evolution of bioluminescence, we will be able to identify new luminescent components quickly and accurately through the genome resource of marine luminous microorganisms.

### Distribution of methane-metabolizing related genomes

Methanogenesis is a strictly anaerobic process in which carbon is used as the electron sink at the absence of oxygen. While biogenic methane is exclusively conducted by methanogens, marine methane can be consumed either aerobically by Proteobacteria or anaerobically by anaerobic methanotrophic archaea (ANME) [83, 84]. Methanogens occupies a wide range of taxonomy with a large proportion belonging to the phylum of Euryarchaeota [27]. These archaea usually use CO2+H2, acetate or other substrates with methyl groups to produce methane. Since one of the key steps in the methanogenic progress is catalyzed by methyl-coenzyme M reductase, its coding gene *mcrA* was widely employed as a marker gene of methanogens. Interestingly, in anaerobic condition, ANMEs and methanogens are genetically close, and both of the microbes possess a typical methanogenesis pathway including *mcr* [85, 86]. ANME cells oxidize methane via a reverse methanogenesis pathway, coupled with reduction of sulphate[27, 87], metal ions[88–90] and nitrate (or nitrite)[91]. On the other hand, in aerobic condition, many reported aerobic methanotrophs belongs to the order Methylococcales of Gamma-proteobacteria or the order Rhizobiales of Alpha-proteobacteria [83, 92]. Methane monooxygenase (MMO) is the key enzyme to perform the oxidization of methane to methanol, and thus the *pmoA* gene which encodes a particulate MMO protein component has been widely used in phylogenetic analyses [93].

In total, 272 genomes were picked out as methane-metabolizing related genomes (MERGs), while 19 genomes were related to aerobic methanotrophs and 253 genomes belong to methanogens and ANMEs (**Fig. 7**). According to the phylogenic trees, the class Methanosarcinia occupies both the most methanogens and ANMEs found in our genome catalogue, while the ANMEs are related to subcluster of ANME-2 and the methanogens are related to those using acetate or substrates with methyl groups. For the subcluster of ANME-1, we found 24 Syntrophoarchaeias, and most of those might be new species (18/24) according to a threshold of ANI > 0.95. As it comes to hydrogenotrophic methanogens, the class of Methanococci and Methanomicrobia contributes the most genomes.

**Figure 7.**
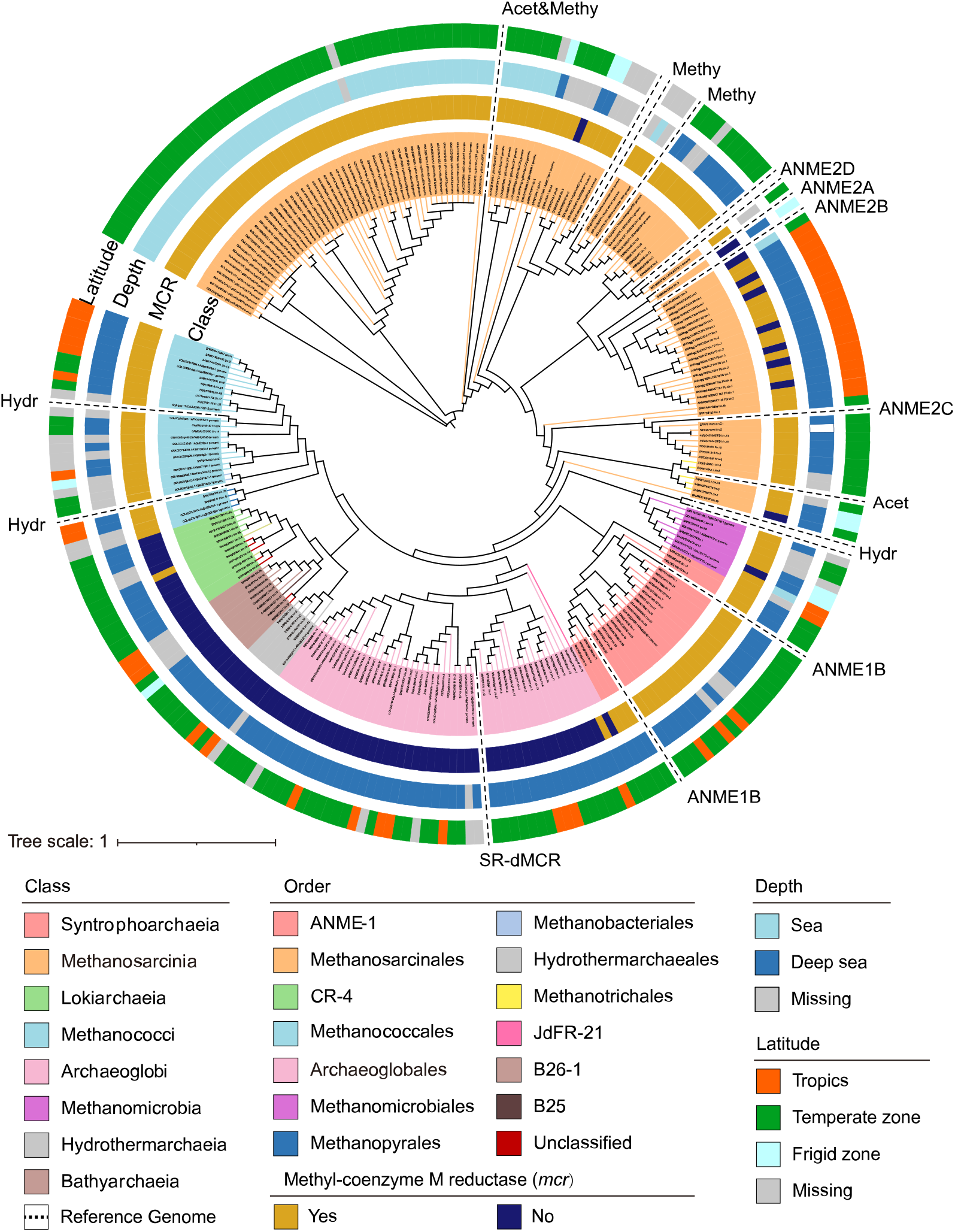
Phylogenetic tree based on archaea genomes of MERGs and reference genomes by GTDB-TK. The classification (Order level and Class level), distribution (Sampling depth and altitude) and the existence of key enzyme coded by *mcr* in methanogenesis pathway were indicated with different colors. Especially, MAGs of unclassified orders were highlighted by red background. Reference genomes were represented by the type of subclusters in white ground (Hydr: hydrogenotrophic methanogenesis, Acet: acetoclastic methanogenesis, Methy: methylotrophic methanogenesis, Acet&Methy: both acetoclastic and methylotrophic methanogenesis, SR&dMCR: a sulfate-reducing archaeon that contained most enzymes for methanogenesis except for *mcr*).

Among all MERGs belongs to archaea, most MERGs were found in deep sea than that of an area of <1000 m in the ocean while other species found >1000 m habitats in sediment, and this pattern is in accordance with the fact that anaerobic conditions is necessary for both methanogenesis or anaerobic oxidation of methane (AOM) (**Fig. 7**)[88, 94, 95]. Interestingly, the distribution pattern in latitude or depth, which is that genomes from same depth group or temperature zones tends to cluster together, indicates that marine prokaryotes with same function, or at least methane-metabolizing related archaea, seem to evolve independently from different geolocations (**Fig. 7**). In addition, there are a large proportion of genomes, which belongs to the lineages of Archaeoglobi, Bathyarchaeia and Hydrothermarchaeia, occupy most enzymes in methanogenesis pathway but lack the key enzyme coding gene of *mcr* (**Fig. 7**). Previous studies have found two Bathyarchaeia genomes which harbored the *mcr* operon [96], as well as several Archaeoglobi genomes [97]. It has been proposed that this *mcr* operon in Bathyarchaeia most likely acquired from euryarchaeotal genomes through horizontal gene transfer [98], or derived from the last common ancestor of Euryarchaeota and Bathyarchaeota [96]. However, as it showed in our results, lineages such as Archaeoglobi and Bathyarchaeia predominantly contain genomes that lack *mcr* operon [98]. Thus, whether these lineages retain *mcr* operon from ancestor or gain mcr through horizontal gene transfer is still inconclusive, and a larger and more systematic dataset would be great help to that. Besides, the absence of *mcr* gene may also reflect the incompleteness of genomes.

## Discussion

The astronomical numbers, incredible diversity, and intense activity of marine prokaryotes have made it a key group in regulating the biosphere, including human being activities, and even the atmosphere, geosphere[3, 5, 6, 99]. Here we analyzed the metagenomic sequencing data of the filtered samples from different oceanic depth layers and the marine sediment samples, host-associated symbiotic samples in each ocean and generated the most integrated marine prokaryotic genome catalogue to date. The resource of 20,671 moderate quality genomes expands the phylogenetic diversity of bacteria and archaea and represents the largest prokaryotic biodiversity in the marine ecosystem. Archaea account for more than 20% of all prokaryotes in seawater, and are the most important microbial group in marine subsurface sediments and most geothermal habitats[27, 100]. In our data, it is currently the largest marine archaeal genome resource dataset, and is the first time to present the phylogenetic tree of global marine archaea containing the most genome level species. Besides, more than 65% phylogenetic diversity was increased of marine prokaryotes, and the diversity increase percentage is consistent with the Earth’s Microbiomes Project [51]. However, inconsistent with the recent studies of microbial diversity[101, 102], two novel candidate Bacteria phyla were detected surprisingly. It indicated that there are still new deep-branching lineages (new phyla or new orders) waiting to be discovered, especially in marine ecosystems. Although we have not been able to collect the whole genomics sequencing data of the entire marine ecosystem, the large-scale marine prokaryotic genome data set currently generated has greatly enhanced our understanding of marine ecosystems and microbial communities. The genome catalogue represents a key step forward towards characterizing the species, functional and secondary metabolite BGCs diversity in marine microbial communities, and will become a valuable resource for future metabolic and genome-centric data mining.

## Method

### Data collection

We compiled all the publicly prokaryotic genomes from NCBI[103] at May 31, 2020. To generated uncluttered genomes, we surveyed the of NCBI, EBI and JGI. In the NCBI database, we screened 55 marine-related Taxonomy ids (**Table S1**). Based on these taxonomy ids, we used NCBI’s E-utilities tool to obtain sample information and sra information, and filtered out non-metagenomic data. Finally, we obtained 26,238 marine metagenomics sample from NCBI public database. In the EBI database, we downloaded the meta data of all classification systems, and then manually screened them according to 27 keywords related to the ocean (**Table S1**), and obtained 5,168 marine metagenomics samples. In the JGI database, we directly used keywords to download relevant sample information, manually corrected it, and finally obtained 82 samples. Because of the data interoperability between different databases, we removed the duplicate data obtained from the three databases and finally got 6265 marine prokaryotic genome samples and 2875 marine metagenomics samples for the downstream analysis.

### Genome binning and quality evaluation

For the metagenomics samples, after filtered low quality, PCR duplication and adapter contamination reads, the clean data of each sample was assembled into contigs by megahit (v1.1) with parameters “--min-count 2 --k-min 33 --k-max 83 --k-step 20”[104]. Subsequently Matabat2 (v2.12.1)[105] module from metawrap (v1.1.5)[106] was used for binning analysis with parameters “−l 1000” to obtain the metagenomics assembled genomes (MAGs).

CheckM (v1.0.12) [107] was used for genome quality evaluation of all public genomes and new MAGs, and the low quality genomes (completeness < 50% or contamination > 10%) was removed. All the moderate genomes (completeness >50% and contamination <10%) were remained and only the substantial genomes (completeness >70% and contamination <10%) were selected for downstream statistics and analysis.

### Species clustering, gene annotation and phylogenetic analyses

The taxonomic annotation of each genome was performed by the Genome Taxonomy Database Toolkit (GTDB-tk, v1.0.2) using the “classify_wf” function and default parameters[108]. To remove redundant genomes, we clustered the total 21,182 substantial genomes at an estimated species level by dRep (v2.6.2)[109] with parameters “-comp 70 -con 10 -pa 0.9 --S_ani 0.95 --cov_thresh 0.3”. The Spearman correlation between genome size and GC content and between the genome features and environmental factors of the major phyla was calculated by R (v3.3.1). All phylogenetic trees were constructed by FastTree (v2.1.10)[110] using the protein sequence alignments produced by GTDB-Tk, and visualized by iTOL (v5.0)[111].

Potential CDS regions of all the microbial genomes, MAGs and metagenome unbinned contigs were predicted by Prokka (v1.14.6)[112], and all predicated CDS sequences were lumped and redundant sequences removed by Linclust [113] to construct a unique gene catalogue for the marine microbiome. The gene sequences of each non-redundant genomes were annotated by KEGG database (v87.0) by Diamond (v0.8.23.85)[114], and secondary-metabolite biosynthetic gene clusters BGCs and regions were identified using antiSMASH (v5.0)[115] with default parameters.

### Methane-metabolizing related genomes detection

Considering the highly shared methane metabolizing pathway either between methanogens and ANMEs or between aerobic methanotrophs, genomes in our genome catalogue which harboring more than 80% of shared KEGG Orthologs of “Methane Metabolism” (Meth-KOs) in several reported species of either methanogens and ANMEs or aerobic methanotrophs (**Table S2**) were picked out as candidates. Candidates were further selected as methane-metabolizing related genomes (MERGs) if one harboring over 50 Meth-KOs. Phylogenic analysis was performed with all MERGs and also the genomes in Table 2.

