## Supplemental figure and tables for "Diversity, function and evolution of marine microbe genomes"

**Supplemental figures and tables.**

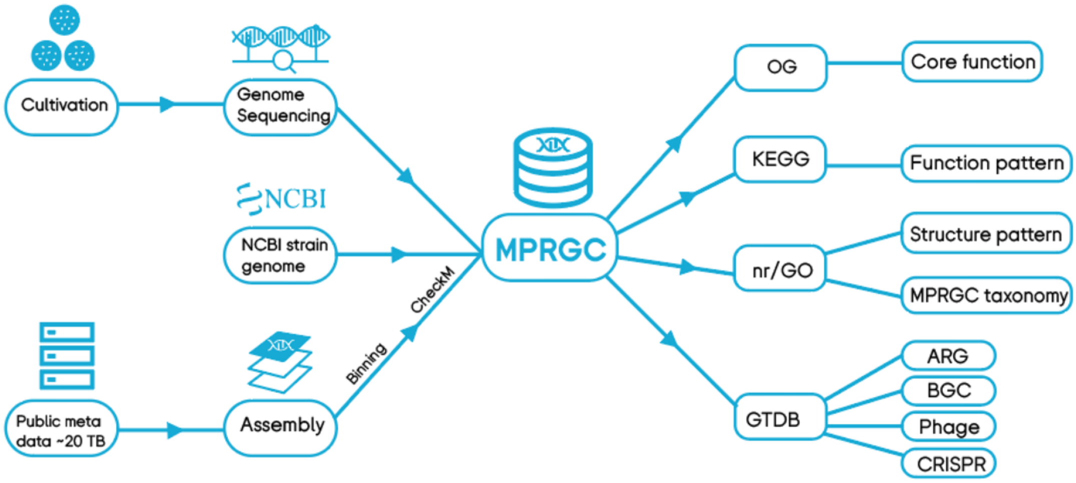


**Fig S1. The overview of the data and methods used to generate the marine prokaryotic reference genome (MPRGC) and protein sequence (MPRPC) catalogs**

**Table S1. The taxonomy names and keywords used in marine prokaryotic data collection**

| NCBI_ taxonomy | NCBI_taxid | EBI/JGI Keywords |
|---|---|---|
| glacier_metagenome | 1651087 | ocean |
| hydrothermal_vent_metagenome | 652676 | seawater |
| invertebrate_gut_metagenome | 1775950 | sea-water |
| invertebrate_metagenome | 1711999 | sea |
| macroalgae_metagenome | 2015907 | marine |
| marine_metagenome | 408172 | bay |
| marine_sediment_metagenome | 412755 | coast |
| microbial_mat_metagenome | 527640 | deepsea |
| rock_metagenome | 1311626 | deep-sea |
| sea_anemone_metagenome | 1825923 | coral |
| sea_squirt_metagenome | 1041057 | shore |
| sea_urchin_metagenome | 1873886 | seashore |
| seagrass_metagenome | 1904484 | microalgal |
| seawater_metagenome | 1561972 | algal |
| sediment_metagenome | 749907 | hydrothermal |
| starfish_metagenome | 2053188 | estuary |
| aquaculture_metagenome | 2714341 | seep |

| | | |
|---|---|---|
| aquatic_metagenome | 1169740 | brine |
| aquifer_metagenome | 1704045 | trench |
| ballast_water_metagenome | 1954210 | mangrove |
| ciliate_metagenome | 1969832 | pelagic |
| ctenophore_metagenome | 1508044 | Atlantic |
| desalination_cell_metagenome | 1983455 | Antarctic |
| eukaryotic_plankton_metagenome | 2315767 | Arctic |
| flotsam_metagenome | 1602165 | Pacific |
| gill_metagenome | 1455666 | phytoplankton |
| hydrozoan_metagenome | 1941281 | Mariana |
| oyster_metagenome | 1541066 | |
| periphyton_metagenome | 1825055 | |
| shrimp_gut_metagenome | 1588881 | |
| zebrafish_metagenome | 1331678 | |
| algae_metagenome | 1300146 | |
| beach_sand_metagenome | 412757 | |
| brine_metagenome | 1981201 | |
| cetacean_metagenome | 1822005 | |
| cold_seep_metagenome | 1583376 | |
| coral_metagenome | 496922 | |
| coral_reef_metagenome | 471232 | |
| crab_metagenome | 1660082 | |
| crustacean_metagenome | 1681198 | |
| dinoflagellate_metagenome | 1579005 | |
| echinoderm_metagenome | 1411990 | |
| estuary_metagenome | 1649191 | |
| fish_gut_metagenome | 1602388 | |
| fish_metagenome | 496924 | |
| jellyfish_metagenome | 1549733 | |
| lagoon_metagenome | 1763544 | |
| mangrove_metagenome | 1284368 | |
| marine_plankton_metagenome | 1874687 | |
| mollusc_metagenome | 1417798 | |
| sand_metagenome | 1671699 | |
| sponge_metagenome | 1163772 | |
| surface_metagenome | 1774230 | |
| tidal_flat_metagenome | 1269027 | |
| whale_fall_metagenome | 412756 | |

**Table S2. Reported Genomes related to methane metabolism**

| Type or Subcluster | NCBI Accession or JGI IMG Numbers | Reference |
|---|---|---|
| **Methanogens** | | |
| Acetoclastic methanogenesis | GCF_000204415.1 | [1] |
| Methylotrophic methanogenesis | GCF_000013725.1 | [1] |
| | GCF_000504205.1 | [1] |
| | GCF_000025865.1 | [1] |
| Hydrogenotrophic methanogenesis | GCF_000013445.1 | [1] |
| | GCF_000302455.1 | [1] |
| | GCF_000016125.1 | [1] |
| | GCF_000327485.1 | [1] |
| Other methanogenesis | GCF_000007345.1 | [1] |
| | GCF_000008665.1 | [1] |
| **Aerobic Methanotrophs** | | |
| Methylococcaceae Type I A | GCA_000685925.1 | [2] |
| | GCA_001312005.1 | [2] |
| | GCA_001644015.1 | [2] |
| | GCA_006788925.1 | [2] |
| Methylococcaceae Type I B | GCA_000008325.1 | [2] |
| | GCA_000427385.1 | [2] |
| | GCA_900155475.1 | [2] |
| Methylothermaceae | GCA_000421465.1 | [3] |
| Beijerinckiaceae | GCA_000427445.1 | [3] |
| | GCA_000385335.1 | [3] |
| Methylocystaceae Type II | GCA_009685195.1 | [3] |
| | GCA_000178815.2 | [3] |
| **ANMEs** | | |
| ANME-1b | GCA_003194425.1 | [4] |
| | GCA_003194435.1 | [4] |
| | GCA_003336485.1 | [4] |
| ANME-2a | IMG_2565956544 | [4] |
| ANME-2b | GCA_002926195.1 | [5] |
| ANME-2c | GCA_003336385.2 | [4] |
| ANME-2d | GCF_000685155.1 | [6] |
| | GCA_001317315.1 | [7] |
